# Anti-phage defence systems are enriched in multidrug-resistant *Pseudomonas aeruginosa*

**DOI:** 10.64898/2026.08.18.745494

**Authors:** Lisa H. Olijslager, Nadiia Pozhydaieva, Stan J. J. Brouns, Antoni P. A. Hendrickx, Pieter-Jan Haas

## Abstract

*Pseudomonas aeruginosa* encodes diverse defence systems against phages and mobile genetic elements, yet their variation across clinical contexts remains unclear. Here, we present a large-scale comparative analysis of the *P. aeruginosa* defensome across both public and clinically highly relevant datasets, including from patients with chronic lung disease and multidrug-resistant isolates. Our analysis shows that while influences on defensome composition are minor, multidrug-resistant isolates encode more defence systems and cystic fibrosis-associated isolates have fewer. Across phylogenetic clusters, defensome size correlates with cluster abundance, suggesting that defence-rich lineages persist more successfully across environments. Lastly, comparative analysis with other *Pseudomonas* species reveals enrichment of anti-plasmid systems in *P. aeruginosa*. Overall, these findings have important implications for phage therapy: multidrug-resistant infections may be more difficult to treat, while cystic-fibrosis-associated isolates may have higher phage susceptibility. This work provides a framework for understanding defensome variation and guiding the decision-making process of phage-based therapies.

## Introduction

Bacteria are continuously challenged by mobile genetic elements (MGEs), such as phages and plasmids (1–6). Over billions of years of coevolution, recurrent phage predation has driven the emergence, diversification, and continual refinement of defence systems that counter MGEs (6–11). Defence systems are frequently encoded in defence islands: genomic loci that cluster multiple systems and facilitate coordinated horizontal gene transfer, leading to novel combinations of defence systems (12–15). Through selective pressure, effective combinations of defence systems are maintained (16–18), resulting in variation in overall defence repertoires (defensomes) between strains depending on ecological and evolutionary contexts (11). However, the clinical factors that shape defensome variation are currently unclear.

*Pseudomonas aeruginosa* is a particularly suitable model for studying defensome variation. It encodes many, diverse defence systems and has a large accessory genome populated by MGEs, enabling rapid adaptation (7,8,13,17–19). Additionally, *P. aeruginosa* is a prime candidate for phage therapy due to rising levels of antimicrobial resistance (20,21). In particular isolates associated with chronic lung disease, such as cystic fibrosis (CF), and carbapenemase-producing *P. aeruginosa* (CPPA) are key targets for therapeutic alternatives. *P. aeruginosa* infections associated with chronic lung disease are often multidrug-resistant and/or embedded in biofilm, which can be difficult for antibiotics to penetrate (22,23). Most CPPA, which are typically resistant to the last-resort antibiotics carbapenems (e.g. meropenem), are multi- or pan-drug resistant (24).

As defensome size correlates with reduced phage susceptibility (8,25), understanding defensome variation in CF-associated and CPPA isolates is critical. Such insights could inform the design and curation of phage collections tailored to specific clinical contexts. Recent studies using large genome datasets have already reported reduced defensomes in cystic fibrosis isolates, but identified different individual defence systems which were involved in this reduction (17,26). This discrepancy could reflect environmental or phylogenetic differences between the used datasets.

In this study, we present a comprehensive analysis of *P. aeruginosa* defensomes using curated clinical datasets of isolates highly relevant to phage therapy: those associated with chronic lung disease (27,28) and those collected for the Dutch surveillance of CPPA. We confirm observations from these datasets using a large public database (the Pseudomonas Genome Database (29)). We analyse defensomes at both the isolate and phylogenetic cluster level, enabling both analysis of real clinical populations as well as in a phylogenetically corrected dataset. We identify differences in defensome size across datasets and show that defensome size correlates with lineage abundance. We assessed how factors such as geographic origin, disease context, and antibiotic resistance relate to defensome variation. Finally, we compared *P. aeruginosa* with other *Pseudomonas* species and found several differences in defensome composition among species. Together, these results show that defensome variation is primarily structured by phylogeny, with additional contributions from clinical context and antibiotic resistance.

## Materials and Methods

### Genomic Databases

This study makes use of three genomic databases. The first database, Chronic lung, originates from an antibiotic-development study for people with chronic lung conditions such as CF (27,28). Isolates in this database originated from respiratory samples from different patients in Australia, the United Kingdom, Spain, and the Netherlands gathered between 2002 and 2016. In total, 411 genomes were successfully sequenced. Of these, 324 were isolates from CF patients. Other isolates were collected from patients with chronic bronchial colonisation, typically due to other confounding chronic lung diseases such as those with bronchiectasis. The second database, Dutch CPPA, is part of the Dutch surveillance for carbapenemase producing *P. aeruginosa*. In this surveillance, Dutch medical microbiology laboratories voluntarily send one *P. aeruginosa* per patient after carbapenemase production and/or presence of a carbapenemase gene is confirmed. In total, 478 *P. aeruginosa* isolates were collected between 2012 and 2025, of which 407 produced carbapenemase (CPPA) and 71 did not. 48 of these were resistant to meropenem (MIC ≥8) in the absence of identified carbapenemase, the remainder were susceptible to meropenem (MIC<8). The third database, Pseudomonas DB, is the Pseudomonas Genome Database (29). This database contained 14,229 genomic sequences at the time of accession (obtained October 2025 from https://www.pseudomonas.com) from different *Pseudomonas* species, 7,981 of which are *P. aeruginosa*. Unless otherwise stated, only the *P. aeruginosa* isolates are considered.

For the Chronic Lung and Pseudomonas DB genomic assemblies could be obtained directly. For the Dutch CPPA strains, Illumina short-read sequencing data was shared, which were assembled using Unicycler version 0.5.1 (47).

The diversity index for each database was calculated by subsampling 33 isolates at random from each of the databases 100 times and calculating the mean of the Shannon diversity index for those isolates.

### Annotation of database metadata

For all three databases, metadata was shared or could be obtained directly. For all, this included geographic location and MLST. For the Pseudomonas DB, the associated country and continent were determined from the location provided using the python package pycountry version 24.6.1.

For the Chronic lung isolates and Pseudomonas DB metadata also included patient disease. For the Pseudomonas DB, disease metadata contained many variations of similar terms. We consolidated similar terms and grouped similar infection types. All metadata terms referring to cystic fibrosis or CF-associated respiratory infections were collapsed into the unified category “cystic fibrosis”. Variants describing bloodstream infections were grouped as “bacteremia”. A broad range of respiratory-related terms (e.g. “pneumonia”, “ventilator-associated pneumonia”, and “respiratory tract infections”) were grouped under “respiratory tract infection”. Terms relevant to urinary infections were grouped as “urinary tract infection”. Finally, wound-related descriptions (e.g., “wound”, “wound discharge”, and “burn wound”) were grouped into “wound infection”.

As the Pseudomonas DB also includes *Pseudomonas* species other than the *aeruginosa* clade, the species was also obtained.

For the Dutch CPPA dataset, additional phenotypic antimicrobial susceptibility test results were obtained by linking the isolates to the database of the national antimicrobial susceptibility surveillance (ISIS-AR). ISIS-AR collects results of routinely performed antibiotic susceptibility testing from bacterial cultures from all healthcare types, to monitor trends in antimicrobial resistance in the Netherlands (48). 65% (313 isolates) from the Dutch CPPA dataset could be linked.

For the Chronic lung and Dutch CPPA dataset, minimum inhibitory concentration (MIC) values or using disk-diffusion zone diameters were converted into categorical susceptibility interpretations for meropenem, ceftazidime, ciprofloxacin, tobramycin, and aztreonam using EUCAST clinical breakpoints (version 15.0, 2025). Isolates were classified as resistant, susceptible at standard dosing (“susceptible, normal exposure”), or susceptible at increased dosing (“susceptible, increased exposure”) according to antibiotic-specific MIC thresholds (49). For analyses concerning antibiotic resistances, susceptible (normal exposure) and susceptible (increased exposure) were both considered susceptible. For the Pseudomonas DB resistance was determined by selecting for “Resistant” or “Susceptible” in the Antibiotic Profile selection filter on Pseudomonas DB specifically for the named antibiotics (accessed: 21-4-2026). For analyses, only isolates were used for which the resistance status was known for all considered antibiotics (meropenem, ceftazidime, ciprofloxacin, tobramycin, and aztreonam (omitted from the Dutch CPPA database due to lack of data)).

### Phylogenetic clustering

Phylogenetic clusters were determined using dRep version 3.6.2 (50). For the Chronic lung database and Dutch CPPA this was done with fastANI (51) as the secondary clustering algorithm. To facilitate dereplication of the large size of the Pseudomonas DB, the accessions were first subdivided by Species based on the metadata. Subsequently, dRep was run with the additional flags multiround_primary_clustering, greedy_secondary_clustering, and primary_chunksize 500. For all, 99.6% was used as a as manually determined sequence identity cutoff value, based on observations of when defence systems started differentiating substantially between genomes.

### Defence system identification

Defence systems were identified using find_prokaryotic_immune_systems version 1.0.0 (34). This algorithm was used to examine and combine the output of DefenseFinder version 1.3.0 (32,52) and PADLOC version 2.0.0 (33), as well as the custom systems from (39,53) in a way designed to easily compare genomic data and without introducing overlap of identified defence systems. PADLOC putative defence systems were not considered.

Genomically clustered data was analysed by collapsing phylogenetic clusters into one “average species”. For example, if a phylogenetic cluster has four genomes, and one has a single representative of a defence system, that cluster will be considered to have 0.25 of that defence system.

### Data visualisation and statistical testing

Defence system totals between databases or subsets of databases were compared using a two-sided Welch’s t-test with Bonferroni correction.

The difference of individual defence system presence within different databases or parameters were compared using a χ^2^-test of independence over the proportion each defence system held within the genome.

Differences in defensome composition were determined by first calculating the proportion of each defence system within the whole defensome and subsequently using a permutational multivariate analysis of variance (PERMANOVA) (35). To this end, raw defence system counts were converted to relative abundances per sample, and samples with zero total abundance were excluded. A centred log-ratio (CLR) transformation was applied to the relative abundance matrix using a pseudocount of 1×10⁻⁶. Aitchison distances were then computed as Euclidean distances in CLR space. Differences were tested using PERMANOVA with 9999 permutations.

Associations between defence system abundance and antibiotic resistance were assessed using multivariable ordinary least squares (OLS) regression.

All statistical tests and data visualisations were performed and generated using the python libraries scipy 1.16.1, statsmodels 0.14.5, skbio 0.7.1.post1, Seaborn version 0.13.2, and matplotlib 3.10.5 in Python version 3.13.2.

## Results

### Defensome comparison of *P. aeruginosa* in clinically relevant datasets

In this study we analysed three *P. aeruginosa* genome databases (**Supplementary data 1**). The first, the Chronic lung database (n = 411), consists of *P. aeruginosa* strains recovered from patients with chronic lung conditions (27,28). The second, Dutch CPPA (n = 478), are the strains sent to the National Institute for Public Health and the environment (RIVM) as part of the Dutch national surveillance for CPPA under suspicion of producing carbapenemase. The third database, the Pseudomonas DB (n = 7,981), are the *P. aeruginosa* isolates in the Pseudomonas Genome Database (29), a large online repository where through a community-driven effort, genomes of different *Pseudomonas* species are collected.

We first assessed database diversity by identifying phylogenetic clusters in the database and using the mean Shannon diversity index of the phylogenetic clusters in each database after subsampling (**Figure 1A-B**). The Dutch CPPA collection was the least diverse, which is consistent with previous epidemiological observations that carbapenemase-producing isolates are often dominated by a limited number of successful lineages at the national scale (30,31). One phylogenetic cluster (largely coinciding with the Multi-Locus Sequence Type 111, of which the prevalence in the Netherlands has been noted before (31)) accounts for 37% of the database. The Chronic Lung and Pseudomonas DB consisted of a larger number of clusters (**Figure 1B**).

**Figure 1:**
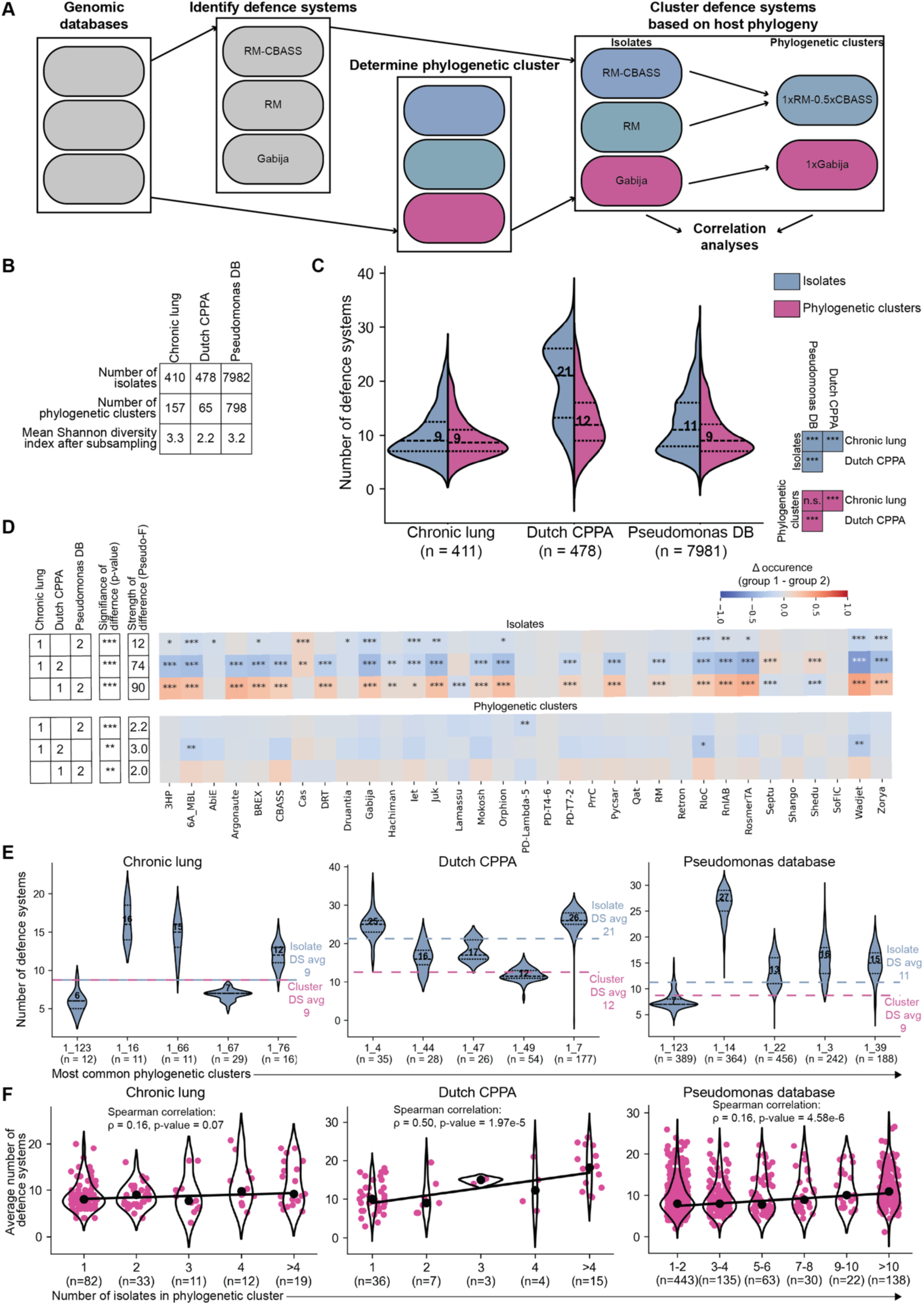
Comparison of defensomes across clinical and reference *Pseudomonas aeruginosa* populations. Systematic comparison of defensome size and distribution of defence systems across three *P. aeruginosa* collections: a cohort of patients with chronic lung infections (Chronic lung), those supplied for the Dutch surveillance of Carbapenemase producing isolates (Dutch CPPA), and one from the Pseudomonas genome database (Pseudomonas DB). **A)** Schematic overview of the bioinformatical workflow. **B)** Summary statistics of each dataset, including number of isolates, number of phylogenetic clusters (defined at 99.6% sequence identity), and diversity (mean Shannon index following subsampling; higher numbers indicate more diversity). **C)** Distribution of defence system counts per genome across datasets, shown for all isolates (blue) and for phylogenetic cluster averages (magenta). Dashed lines indicate medians; isolates numbers (n) are shown. Statistical comparisons were performed using a Welch’s t-test. **D)** Differences in overall defence composition between datasets assessed by PERMANOVA. Heatmap shows system-specific differences, represented as proportional enrichment between pairwise comparisons (where 2 was compared with number 1), with significance determined by χ² tests. Only systems contributing ≥15% of the average defensome in at least one dataset are shown. **E)** Defensome distributions for the five most abundant phylogenetic clusters, with their means (black dashed line) compared to overall isolate-(blue dashed line) and cluster-level (magenta dashed line) means. **F)** Relationship between cluster prevalence and defensome size. Phylogenetic clusters were binned by abundance, and their average number of defence systems is shown. A positive association is observed, with more abundant clusters encoding larger defensomes. Violin plots show distributions within bins, with individual datapoints represented as magenta markers and medians indicated with black markers. The trendline is fitted to bin medians. Number of phylogenetic clusters in each bin are indicated (n). Spearman’s ρ (based on all datapoints) and corresponding p-values quantify the strength and significance of the association. *** denotes a p-value < 0.0005, ** a p-value < 0.005, * a p-value < 0.05, and n.s. denotes not significant.

We next quantified defence system repertoires across all genomes using a combined DefenseFinder (32) and PADLOC (33) approach (34) (**Figure 1A**; **Supplementary data 2**). Defensome size differed significantly between all three databases at the isolate level (Welch’s t-test p-value < 0.005; **Figure 1C**). To assess the contribution of phylogeny, we analysed the distribution of average defensome sizes of phylogenetic clusters (**Figure 1A, C**). After aggregation at the cluster level, the Dutch CPPA collection still differed significantly from the other two datasets (p-value < 0.005), whereas the Chronic Lung and Pseudomonas DB were not statistically significantly different.

We next compared the composition of the defensomes (**Figure 1D**). To compare the differences in the total composition of the defensomes independent of defensome size we used a PERMANOVA test, which is designed to find differences in ecological multivariate data, like microbiome compositions (35). This analysis revealed statistically significant differences in overall composition across datasets at both the isolate and phylogenetic cluster levels (PERMANOVA p < 0.005). However, effect sizes at the cluster level were small (Pseudo-F < 3.0), indicating that compositional differences largely reflect shifts in lineage abundance rather than large systematic differences in defence system usage. Consistent with this, few individual defence systems showed significant enrichment or depletion when analyses were restricted to phylogenetic clusters (**Figure 1D**). These findings indicate that, though there are differences in defensome size between the databases, these differences are not driven by specific individual defence systems.

Together, these results indicate that *P. aeruginosa* populations from different clinical and surveillance contexts differ substantially in defensome size and composition. Much of this variation is explained by differences in phylogenetic structure, highlighting the importance of accounting for lineage composition when comparing defence system repertoires across datasets. However, even after correcting for this, *P. aeruginosa* in the Chronic Lung and Dutch CPPA database still differed significantly in defensome size, indicating there are epidemiological parameters that influence the defensomes of these different patient population.

### Phylogenetic cluster prevalence correlates with defensome size

We observed that several abundant phylogenetic clusters encode elevated numbers of defence systems (**Figure 1E**). We therefore tested if there a general association between phylogenetic cluster abundance and defensome size using a Spearman correlation analysis (**Figure 1F**). Across all three databases, we observed a positive association. The strength of this association varied between datasets: it was weak and not statistically significant (ρ > 0.2; p-value > 0.05) in the Chronic Lung database, weak but statistically significant in the Pseudomonas Genome Database (ρ < 0.2, p-value < 0.0005), and strongest in the Dutch CPPA dataset, where a moderate correlation was observed (0.2 < ρ < 0.6, p-value < 0.0005).

As disease context differs substantially between these collections, we investigated if the strength of this correlation depended on infection type (**Supplementary Figure S1**). A positive correlation (Spearman correlation p-value < 0.05) between cluster size and defensome size was observed for all disease contexts. However, the association was notably weaker for cystic fibrosis–associated isolates (ρ < 0.2) than for bloodstream, urinary tract, intra-abdominal, or respiratory tract infections (0.2 < ρ < 0.6). Importantly, respiratory tract infections showed the strongest correlation despite sharing a similar infection site with cystic fibrosis, indicating that chronic infection dynamics rather than the anatomical location likely underlie this difference. These results suggest that lineages encoding larger defence repertoires are more likely to be over-represented in genomic datasets, but that this advantage is reduced in chronic lung infections, where selective pressures may differ.

Overall, these results indicate that defensome size is linked to the epidemiological success of phylogenetic lineages, although the strength of this association depends strongly on clinical context.

### Clinical contexts determine defensome variation in *P. aeruginosa*

The two clinical genome collections analysed in this study, the Chronic Lung and Dutch CPPA dataset, differed in defensome sizes (**Figure 1C**). Because these collections represent clinical contexts highly relevant to phage therapy, we assessed which epidemiological parameters underly these differences.

One distinction between the datasets is their geographical composition. Whereas the Dutch CPPA collection is confined to the Netherlands (although some patients were recently hospitalised abroad), the Chronic Lung dataset includes isolates from multiple European countries and Australia, and samples of the Pseudomonas DB originate from across the world. We compared defensome size and composition at three levels: regionally (using the metadata of the Dutch CPPA dataset for several provinces in the Netherlands; **Supplementary Figure S3**), nationally (for several countries with different antibiotic usage profiles in Europe from the Pseudomonas DB; **Supplementary Figure S4**), and continentally (based on data from the Pseudomonas DB; **Supplementary Figure S5**). Across comparison levels, defensome size and composition differed significantly between geographic regions when isolates were compared. However, these differences were largely abolished after accounting for phylogenetic structure, indicating that geographic variation primarily reflects lineage composition.

Another difference between the two clinical collections is patient disease context. The Chronic Lung dataset consists of isolates from patients with chronic lung conditions, whereas the Dutch CPPA dataset represents a broader clinical spectrum, with only 70 samples originating from sputum (15% of database), and others ranging from colonisation screening swabs (e.g. rectal and perineal swabs) to bloodstream infections. We therefore analysed defensome variation across disease contexts (**Figure 2A**). Consistent with previous reports (17,26), *P. aeruginosa* isolates and phylogenetic clusters from cystic fibrosis patients encoded significantly fewer defence systems than other clinical isolates (defence system median: 8 and 10-16, respectively; Welch’s t-test p-value < 0.005; **Figure 2A**). In a few cases, this reduction was accompanied by a modest but statistically significant shift in defensome composition (PERMANOVA Pseudo-F 1.9-2.5, p-value < 0.005; **Supplementary Figure S5A**).

**Figure 2:**
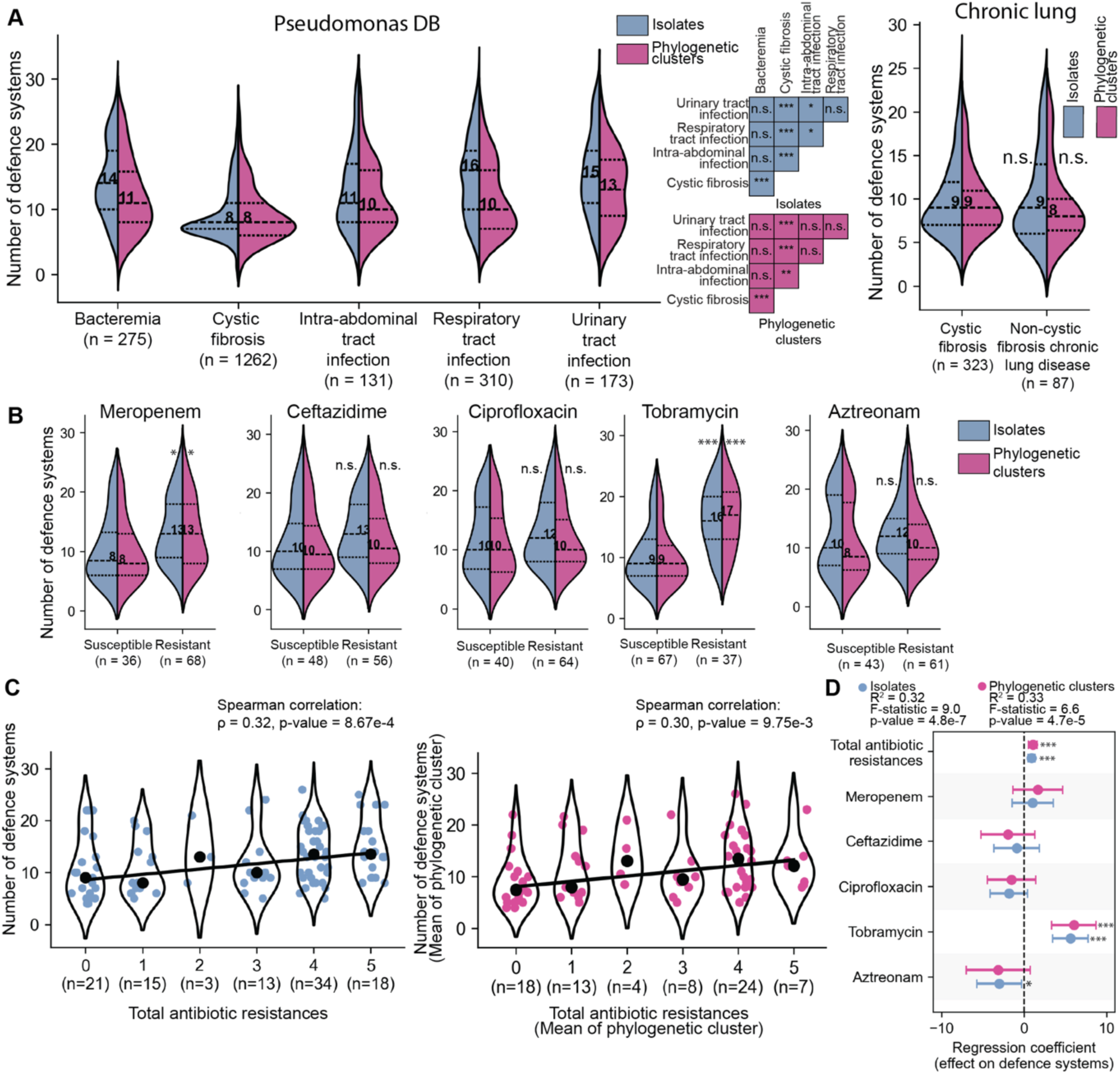
Disease context and antibiotic resistance are associated with variation in *Pseudomonas aeruginosa* defensome size. Evaluation of how defence system abundance varies across disease context and antibiotic resistance phenotypes. **A)** Distribution of defence system counts stratified by patient disease indication, shown at both the isolate level (blue) and as phylogenetic cluster averages (magenta) for the Pseudomonas DB (left) and Chronic lung database (right; subdivided in cystic fibrosis-associated and non-cystic fibrosis-associated). Medians are indicated (black dashed line), and statistical significance was assessed using Welch’s t-test for isolate-level (blue) and cluster-level (magenta) comparisons. **B)** Association between specific antibiotic resistances and defence system abundance, excluding cystic fibrosis-associated isolates. Isolates were classified as resistant or susceptible to five antibiotics spanning distinct classes, shown at both the isolate level (blue) and as phylogenetic cluster averages (magenta) for the Pseudomonas DB. Medians are indicated (black dashed line); statistical significance was determined using a Welch’s t-test. **C)** Relationship between cumulative resistance to antibiotics and defence system abundance. The number of antibiotics to which an isolate (left) or phylogenetic cluster (right; average number of resistances) is resistant is plotted against its corresponding (average) defensome size. Distributions are shown as violin plots, with individual data points overlaid (blue or magenta markers of isolates and phylogenetic clusters, respectively) and medians indicated (black markers). Trendlines were fitted to bin medians, and Spearman’s ρ and associated p-values are shown to indicate the strength and significance of the association, respectively. **D)** Multivariable regression analysis quantifying the association between antibiotic resistance and defence system abundance. Forest plot shows regression coefficients with 95% confidence intervals for multidrug resistance overall and for individual antibiotics. Positive coefficients indicate increased defence system abundance associated with resistance. Models were fitted at both the isolate level (blue) and phylogenetic cluster level (magenta). *** denotes a p-value < 0.0005, ** a p-value < 0.005, * a p-value < 0.05, and n.s. denotes not significant.

Comparison within the Chronic Lung dataset showed no significant differences between cystic fibrosis and non–cystic fibrosis isolates, indicating that the reduced defensome size is a broader feature of chronic lung-associated *P. aeruginosa* rather than being a unique for cystic fibrosis isolates (**Figure 2B**).

The Dutch CPPA dataset, which largely consists of carbapenemase-producing *P. aeruginosa*, exhibits significantly larger defensomes than the other collections (**Figure 1C**). To test whether this enrichment was linked to meropenem resistance (a common example of an antibiotic in the carbapenem class) and/or multidrug resistance (defined as cumulative resistance to different antibiotic classes) in general, we compared defensome size between resistant and susceptible isolates in the Chronic lung (**Supplementary figure S6A**), Dutch CPPA (**Supplementary figure S7A**), and Pseudomonas DB (**Figure 2B**, cystic-fibrosis-associated samples excluded). This analysis included meropenem (a carbapenem) as well as ceftazidime, ciprofloxacin, tobramycin, and aztreonam (omitted for Dutch CPPA due to lack of data). These antibiotics are representatives of five antibiotics classes that are commonly used for treating *P. aeruginosa* infections. While in the Chronic lung dataset limited effects were seen (**Supplementary figure S6A**), likely due to the strong confounding influence of chronic lung disease, significantly more defence systems were found in both the meropenem-resistant (Pseudomonas DB only; median number of defence systems: 8 (susceptible) and 13 (resistant, Welch’s T-test p-value < 0.05; **Figure 2B**) and tobramycin resistant populations (Pseudomonas DB and Dutch CPPA database; median number of defence systems: 9 (susceptible) and 16-17 (resistant), and 9-10 (susceptible) and 14-20 (resistant), respectively, Welch’s T-test p-value < 0.0005; **Figure 2B**, **Supplementary S7A**). There was little to no difference in the compositions of the antibiotic resistant and susceptible populations (**Supplementary figure S5B, S6D, S7D**). We also observed a moderate correlation between the number of antibiotic resistances and defensome size in both the Pseudomonas DB and Dutch CPPA databases (Spearman correlation 0.2 < ρ < 0.6 (except for Dutch CPPA isolates (Spearman correlation ρ < 0.2)), p-value < 0.05; **Figure 2C, Supplementary figure S7B).**

As multidrug resistance and resistances to specific antibiotic are correlated, we performed multivariable regression analysis to distinguish what the effect of each individual antibiotic and multidrug resistance in general is. We found highly statistically significant correlation of moderate strength (isolates: R^2^ = 0.20-0.33 and p-value < 0.05). Out of the modelled parameters (**Figure 2E**), multidrug resistance in general (i.e. cumulative resistance to different antibiotic classes; regression coefficient: 0.89-1.8) had a small effect and tobramycin resistance (regression coefficient: 4.9-6.0, p-value < 0.0005, with exception of Dutch CPPA clusters (p-value = 0.06)) had a pronounced effect on the total number of defence systems.

Together, these analyses show that the difference in defensome size between the two curated clinical collections may largely be explained by opposing clinical pressures. Isolates associated with chronic lung infections, particularly cystic fibrosis, encode smaller defensomes, whereas multidrug resistance in general and Tobramycin resistance in particular are positively correlated with defensome size. Importantly, in both cases, most of the observed variation in defensome size could be attributed to differences in the abundance of phylogenetic lineages.

### Defensome composition varies between *Pseudomonas* species

We next compared defensome composition across different *Pseudomonas* species to place these patterns in a broader, non-clinical context. We analysed several *Pseudomonas* species that were well-represented in the Pseudomonas DB (*P. ffuorescens*, *P. putida*, *P. sp.*, *P. syringae*). These species occupy diverse environmental niches and are less or non-pathogenic, providing a useful contrast to *P. aeruginosa*, which is a highly adapted opportunistic pathogen.

We first examined the number of defence systems of each of the *Pseudomonas* species (**Figure 3A**). Defensome size differed modestly (median defence system number: 9-11 vs 9-11 in *P. aeruginosa*) with the exception of *P. syringae*, which differed to a larger degree (median defence system number: 15-16).

**Figure 3:**
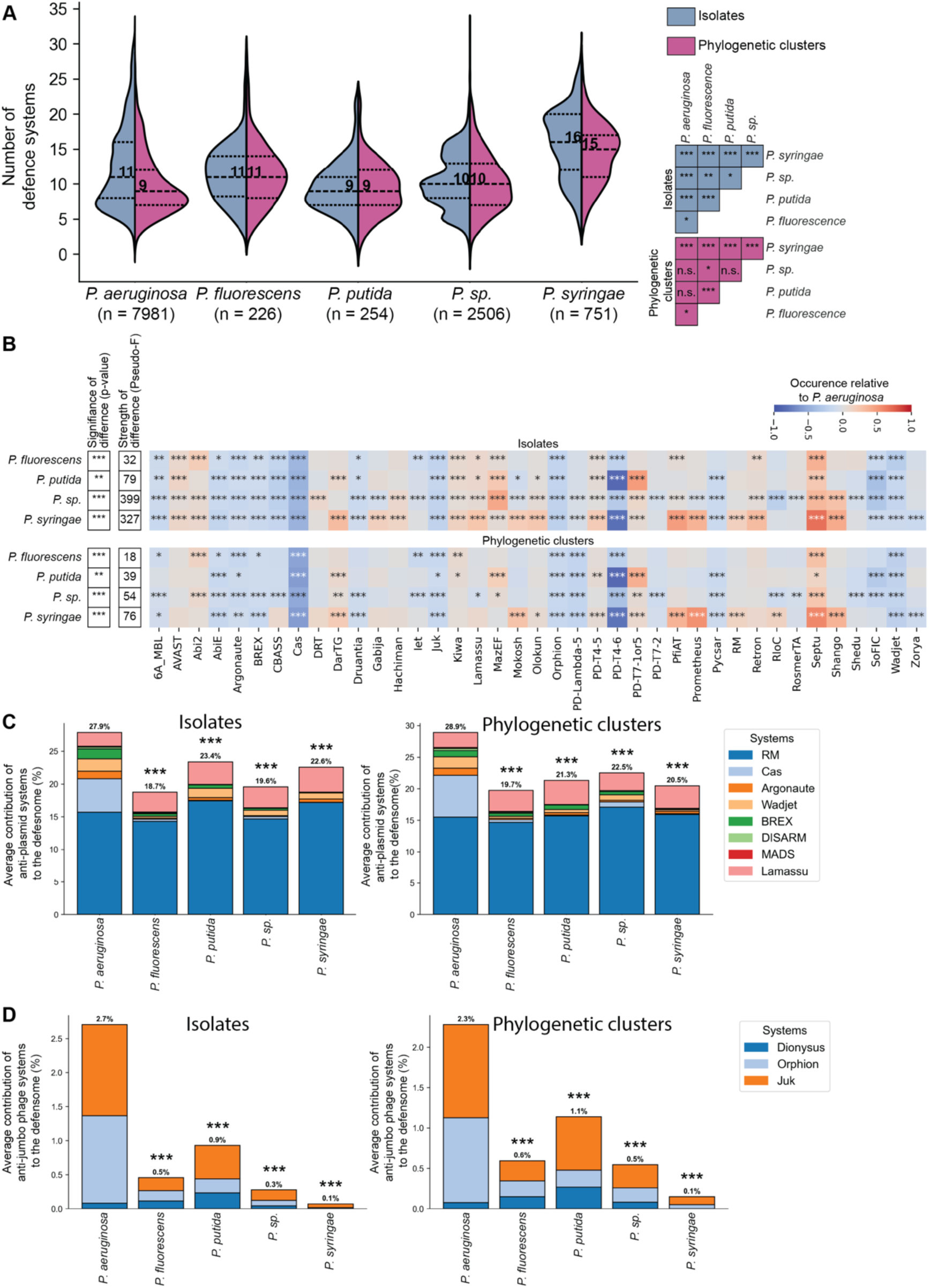
Defence repertoires differ across *Pseudomonas* species, with enrichment of anti-plasmid and anti-jumbo phage systems in *Pseudomonas aeruginosa*. Comparison of defence system abundance and composition across multiple *Pseudomonas* species represented in the Pseudomonas Genome Database. **A)** Distribution of defence system counts across species, shown at both the isolate level and phylogenetic cluster level. Medians are indicated (black dashed line), and statistical significance was assessed using Welch’s t-test for isolate-level (blue) and cluster-level (magenta) comparisons. **B)** Pairwise comparisons of defence system composition between species relative to *P. aeruginosa*. Differences in overall composition were assessed using PERMANOVA. The heatmap shows system-specific proportional differences in prevalence, with significance determined by χ² tests. Only systems contributing ≥15% of the average defensome in at least one species are shown. **C-D)** Contribution of anti-plasmid (**C**) and anti-jumbo phage (**D**) defence systems to total defensomes across species, shown as stacked bar plots. Both individual systems and aggregated categories are displayed. Statistical significance of differences relative to *P. aeruginosa* was assessed using Welch’s t-test. *** denotes a p-value < 0.0005, ** a p-value < 0.005, * a p-value < 0.05, and n.s. denotes not significant.

In contrast, defensome composition differed strongly between *P. aeruginosa* and other *Pseudomonas* species at both the isolate and phylogenetic cluster level (**Figure 3B**). Several systems were specifically enriched or depleted (**Figure 3B**). Notably, of these, multiple systems that have been shown to defend against plasmids (Argonaute, Cas, and Wadjet (36,37)) and jumbo phages (Juk (38) and Ophion (39)) were enriched in *P. aeruginosa*.

To assess broader functional trends, we grouped defence systems by activity against plasmid (in addition to the previous: RM, BREX, DISARM, Lamassu, and MADS (40–43); **Figure 3C**) and jumbo-phages (in addition to the previous: Dionysus (39); **Figure 3D**). *P. aeruginosa* encoded a significantly higher proportion of these systems at both the isolate and cluster level. Anti-plasmid systems contributed approximately 1.2–1.5-fold more to the defensome, while anti-jumbo-phage systems were enriched by 3- to 27-fold, depending on the level of analysis (Welch’s t-test p-value < 0.005). This enrichment, however, was not driven by all systems within each category (**Supplementary Figure S8**; **Figure 3D**). RM systems contributed similarly across species, and Lamassu systems were more prevalent in some non-*aeruginosa* species, possibly reflecting the potential of both these systems to target invaders beyond plasmids (44). Among anti-jumbo-phage systems, Juk and Ophion were consistently enriched in *P. aeruginosa*, whereas Dionysus showed no clear species-specific trend. Seeing as Juk and Ophion often colocalise (81% of the time in our dataset) (39), it is unclear whether this reflects increased evolutionary pressure by jumbo phages, or if there is an alternative explanation for the specific enrichment of Juk and Ophion.

Finally, we investigated whether the link between phylogenetic cluster size and defensome size observed in *P. aeruginosa* (**Figure 1F**) extended to other *Pseudomonas* species. Weak positive correlations were detected for *P. putida* and *P. syringae*, whereas no significant association was observed for *P. fluorescens* or *P. sp.* (**Supplementary Figure SG**).

Together, these results demonstrate that *P. aeruginosa* differs markedly from other *Pseudomonas* species in the functional composition of its defence arsenal, particularly through the enrichment of systems targeting plasmids.

## Discussion

*P. aeruginosa* carry a diverse and expansive set of defence systems (7,8,13,17,18). In this study we performed a large-scale comparative analysis on how defence systems vary based on several clinical parameter. We did this by comparing two datasets consisting of clinical isolates relevant for phage-therapy and using that as a guideline to confirm observations in a large database. By combining both big data with curated patient datasets, we identified multiple correlations that were both statistically robust and biologically meaningful. Analysing both isolates and phylogenetic clusters allowed us to distinguish phylogenetic effects (which were substantial) from epidemiological associations.

Defensome size is associated with two key factors. First, *P. aeruginosa* associated with CF (and likely chronic lung infections in general) encode fewer defence systems. Second, multidrug resistance is positively associated with defensome size. Additionally, defence-rich phylogenetic clusters are more prevalent across datasets. The strength of this correlation depends strongly on disease type and was minor for CF patients and strong in the Dutch CPPA dataset. No individual systems consistently drove changes in defensome size, instead broad shifts across multiple systems were responsible. These findings suggest that clinical context shapes the selective pressures acting on defence repertoires.

As postulated before (17,26), we also suggest that CF-associated *P. aeruginosa* are highly specialised to their environment, meaning they likely encounter fewer or a lower diversity of phages. For the correlation between defensome size and multidrug resistance, an observation which, to the best of our knowledge, has never been observed before, we postulate three possible explanations. First, it is possible that *P. aeruginosa* are more likely to cause more severe infection (leading to an increased likelihood of being sequenced) if they have more defence systems because bacterial populations are not being curbed by phages in the human host. Second, it is possible that *P. aeruginosa* are more likely to encounter (new) phages when adapting to a new environment, for example while colonising a new host or when moving between hosts, meaning that *P. aeruginosa* enriched in defence systems are more likely to successfully spread from patient to patient. This would also explain why *P. aeruginosa* associated with CF, which rarely encounter new environments, don’t have the same evolutionary pressure to incorporate or maintain a wide variety of defence systems. Third, it is possible that since both antibiotic resistance and defence systems mainly spread through mobile genetic elements (1,5), that general enrichment of mobile genetic elements is responsible for both enrichments. This could also explain the particularly strong correlation between Tobramycin resistance and defence systems, if these are mobilised through similar mobile genetic elements.

Differences in defensome composition between *P. aeruginosa* groups were largely tied to phylogeny. However, when we compared *P. aeruginosa* with other *Pseudomonas* species that are less or non-pathogenic, we found substantial differences in defensome composition, especially where anti-plasmid systems are concerned. This might seem counter-intuitive, as *P. aeruginosa* is known to evolve quickly to counter antibiotics (45), and plasmids are typical carriers of antibiotic resistance genes (46). However, plasmids can also pose a significant metabolic burden and antibiotic resistance can also be located on other mobile genetic elements (44). We therefore hypothesise that *P. aeruginosa* diminishes plasmids to reduce metabolic burden so they can persist in a hostile environment, similar to what has already been shown for *Vibrio cholera* (41).

Our findings have direct implications for phage therapy. Although our results suggest that phage panels developed in one geographic region may still retain broad effectiveness elsewhere, regional differences in lineage composition may still influence treatment outcomes.

Next, we found more anti-phage defence systems in multidrug-resistant isolates, which are prime candidates for phage therapy, which may complicate the development of effective phages or phage cocktails. As a practical example, the largest phylogenetic cluster in the Dutch CPPA database (comprising 37% of the entire Dutch CPPA database), has an average of 26 defence systems (**Figure 1E**). Previous research has shown that *P. aeruginosa* with sizable defensomes were often omni- or pan resistant to phages (7). Additionally, even within phylogenetic clusters, there were still frequently differences in some defence systems (**Figure 1E**), likely caused by the evolutionary-arms race between phages and bacteria, causing these systems to be amongst the first to change (9). This means that even after creating a specialised phage cocktail for a prevalent strain like this, defensome diversity within the cluster could render the phage cocktail ineffective.

However, reduced defence repertoires in CF-associated isolates, which are also interesting candidates for phage therapy, may mean increased susceptibility to phage therapy.

Taken together, this work provides a framework for understanding the epidemiology and evolution of the *P. aeruginosa* defensome. The links found between defensome size, phylogeny, clinical context, and antimicrobial resistance have important implications for phage therapy and our broader understanding of bacterial defence system evolution.

## Supplementary Data

Supplementary Data are available.

## Data availability

All data necessary to evaluate the conclusions in the paper are present in the paper or the Supplementary Materials. The Dutch CPPA database is available at Genbank bioproject PRJNA1489403.

## Author contributions

L.H.O., P.J.H., and S.J.J.B. conceived the project. L.H.O. wrote scripts, generated, and analysed all data, performed statistical analyses and made figures under supervision of P.J.H. L.H.O. wrote the manuscript with input from A.P.A.H., N.P., and S.J.J.B.. Funding was acquired by P.J.H. and S.J.J.B.

## Acknowledgement

We are grateful to Ingrid Witt and Desi Fierlier for insightful discussions and valuable suggestions that contributed to this work. We also thank the members of the Medical Microbiology laboratory at UMC Utrecht and Brouns lab at the Delft University of Technology for constructive discussions and feedback throughout the project. Finally, we would like to extend our appreciation to the ISIS-AR project team, ISIS-AR study group, and the participating medical microbiology laboratories for the phenotypic data of the Dutch CPPA strains.

