## Supplementary figures for "Anti-phage defence systems are enriched in multidrug-resistant *Pseudomonas aeruginosa*"

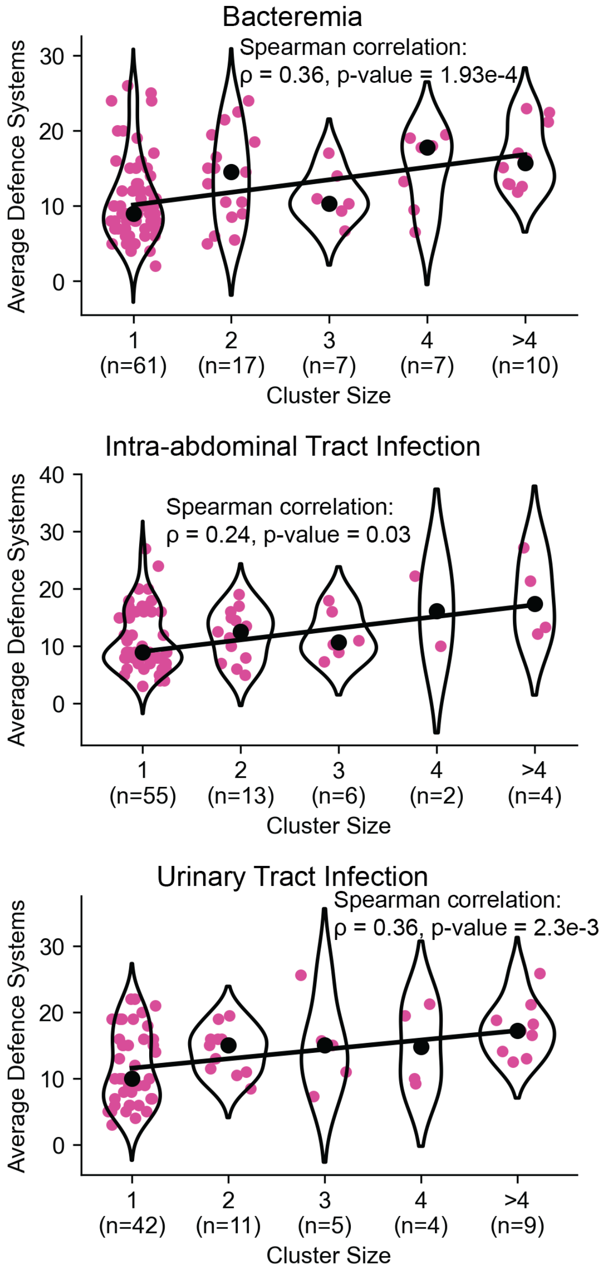


**S1:** Relationship between the number of isolates of phylogenetic clusters and the average number of defence systems per cluster for the *P. aeruginosa* genomes in the Pseudomonas Genome Database for different patient diseases. Clusters were grouped in bins based on the number of isolates of that cluster. Violin plots show the distribution of the different average defence systems of a cluster per bin, individual datapoints are represented in magenta and the median of each bin is indicated with a black marker. The linear trendline is fitted through the medians of these bins. The Spearman correlation coefficient (ρ) and p-values are reported to indicate the strength and significance of the association, respectively.

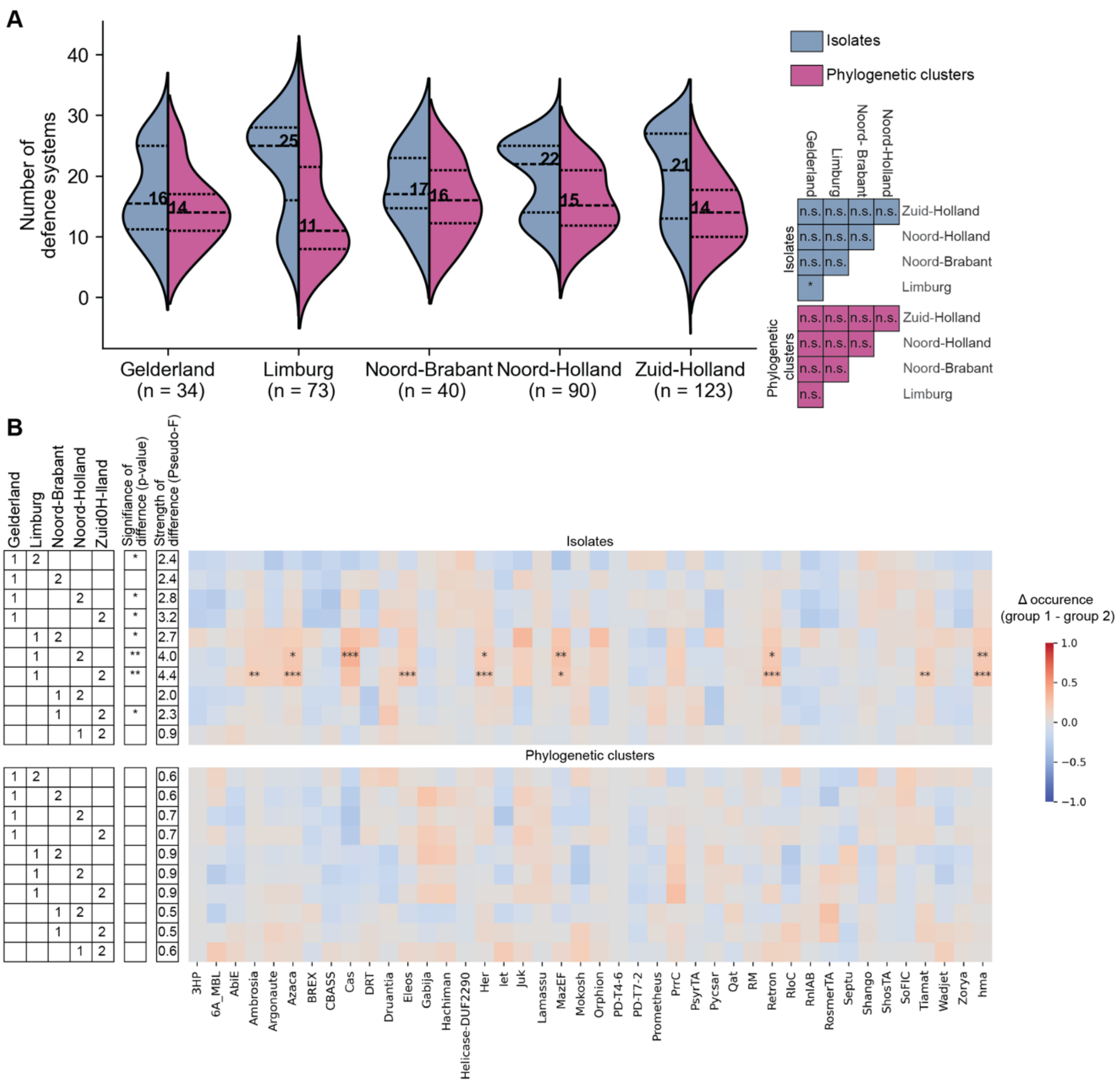


**S2**: Evaluation of the defence systems in different provinces of the Netherlands in the Dutch CPPA database. Only provinces with a minimum of 30 isolates were considered **A)** Violin plot showing the distribution of defence systems across isolates or by averaging over each phylogenetic cluster. Medians are indicated. Statistical significance of the differences between the databases were calculated using a Welch's T-test for both all isolates (blue) and the phylogenetic clusters (magenta). **B)** Comparisons of the defence system composition. Differences in the total composition were determined using a PERMANOVA test. Heatmap shows the comparisons of individual defence systems between the different provinces, comparing the proportional difference of how frequently a defence system is found and the statistical significance, based on the p-value of a $\chi^2$-test of independence. Only systems composing a minimum average of 15% of the total defensome in one of the datasets were considered. *** denotes a p-value < 0.0005, ** a p-value < 0.005, * a p-value < 0.05, and n.s. denotes not significant.

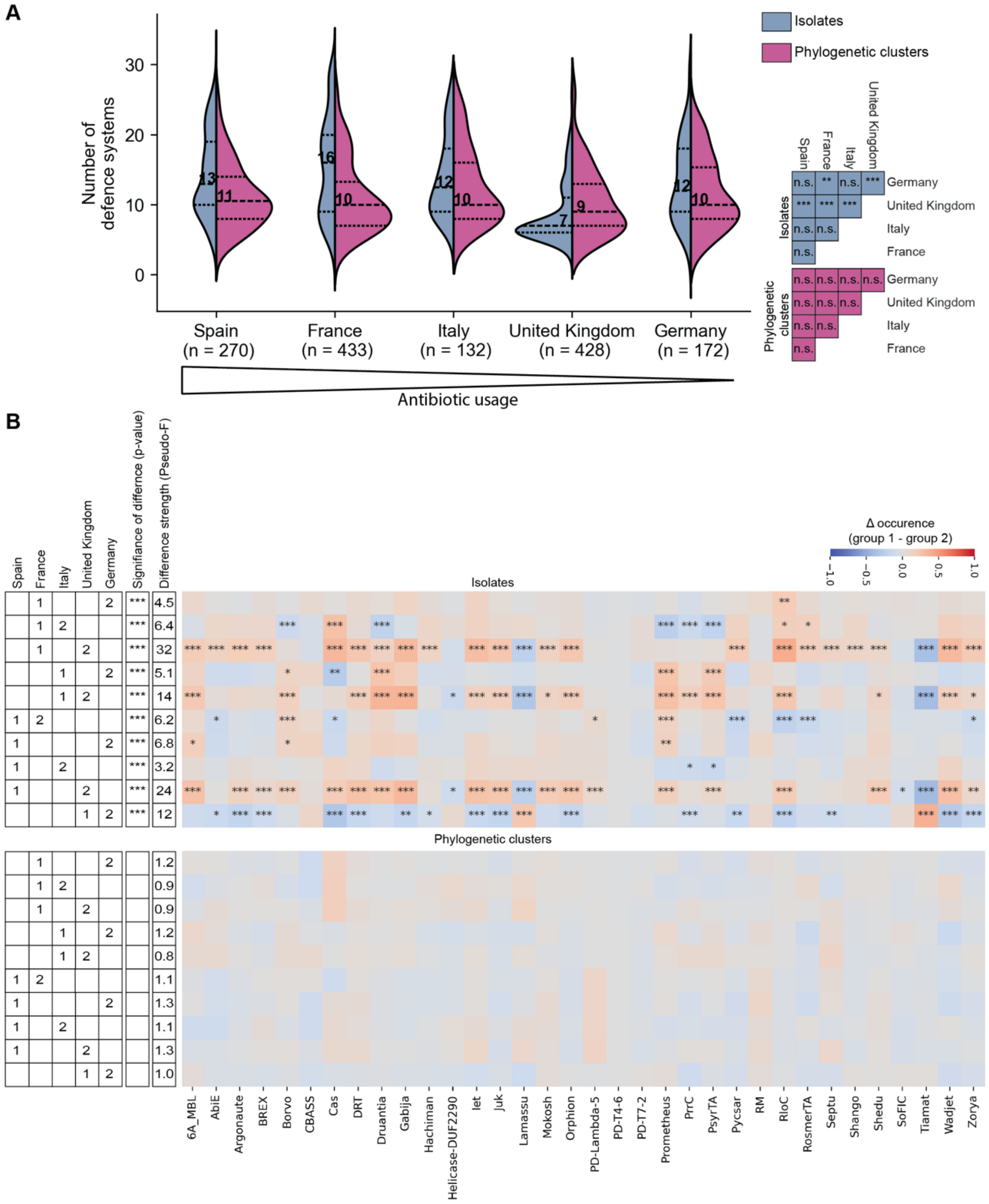


**S3**: Evaluation of the defence systems in *P. aeruginosa* in the Pseudomonas genome database from selected European countries with different antibiotic usages. Countries are ordered by antibiotic usage from high (left) to low (right) (*43*). **A)** Violin plot showing the distribution of defence systems for different countries both across isolates or by averaging over each phylogenetic cluster. Medians are indicated. Statistical significance of the differences between the countries were calculated using a Welch's T-test for both all isolates (blue) and the phylogenetic clusters (magenta). **B)** Comparisons of the defence system composition. Differences in the total composition were determined using a PERMANOVA test. Heatmap shows the comparisons of individual defence systems between the different countries, comparing the proportional difference of how frequently a defence system is found and the statistical significance, based on the p-value of a $\chi^2$-test of independence. Only systems composing a minimum average of 15% of the total defensome in one of the datasets were considered. *** denotes a p-value < 0.0005, ** a p-value < 0.005, * a p-value < 0.05, and n.s. denotes not significant.

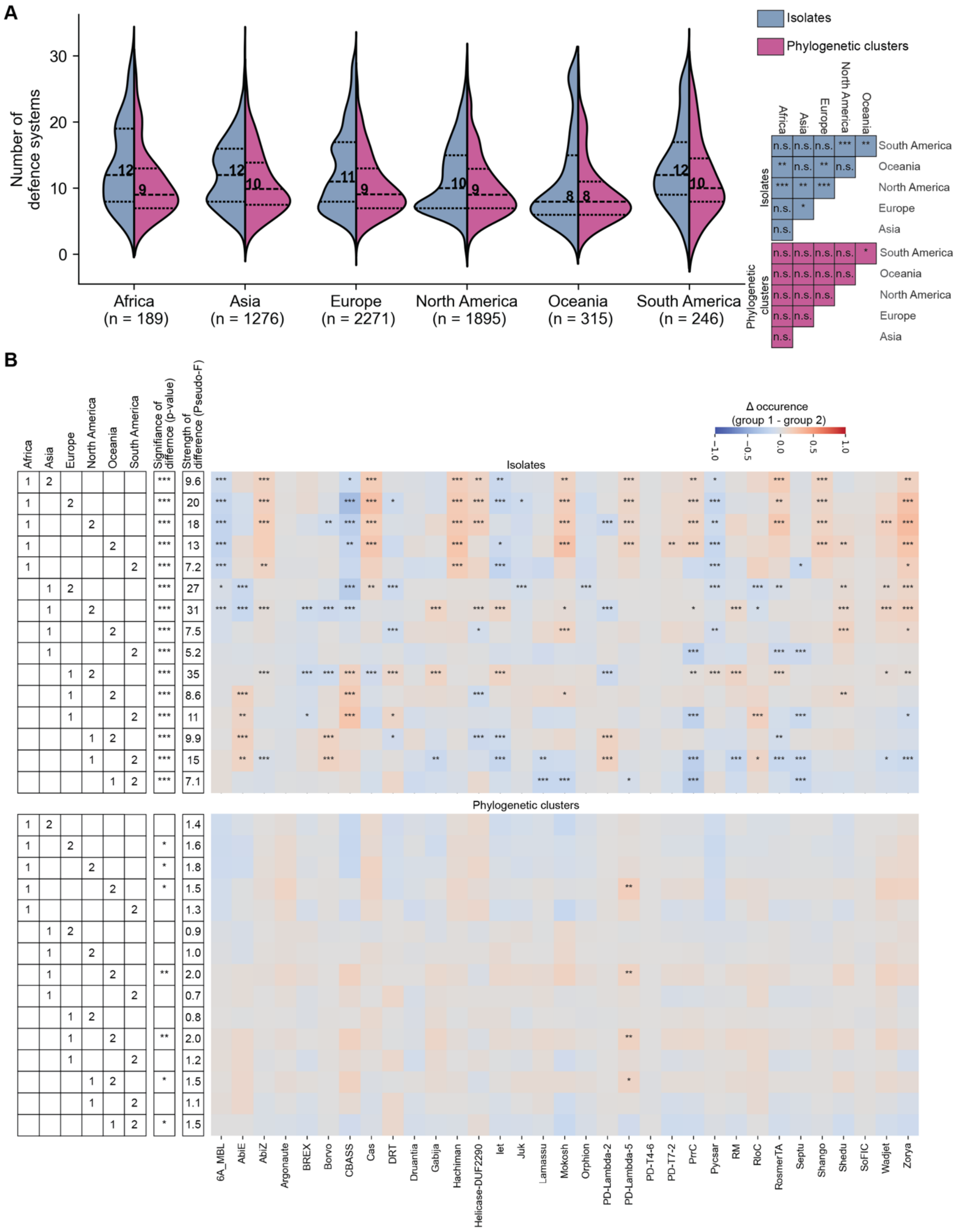


**S4:** Evaluation of the defence systems in *P. aeruginosa* in the Pseudomonas genome database for each continent. **A)** Violin plot showing the distribution of defence systems for different continent both across isolates or by averaging over each phylogenetic cluster. Medians are indicated. Statistical significance of the differences between the databases were calculated using a Welch's T-test for both all isolates (blue) and the phylogenetic clusters (magenta). **B)** Comparisons of the defence system composition for different continents. Differences in the total composition were determined using a PERMANOVA test. Heatmap shows the comparisons of individual defence systems between the different continents, comparing the proportional difference of how frequently a defence system is found and the statistical significance, based on the p-value of a $\chi^2$-test of independence. Only systems composing a minimum average of 15% of the total defensome in one of the datasets were considered. *** denotes a p-value < 0.0005, ** a p-value < 0.005, * a p-value < 0.05, and n.s. denotes not significant.

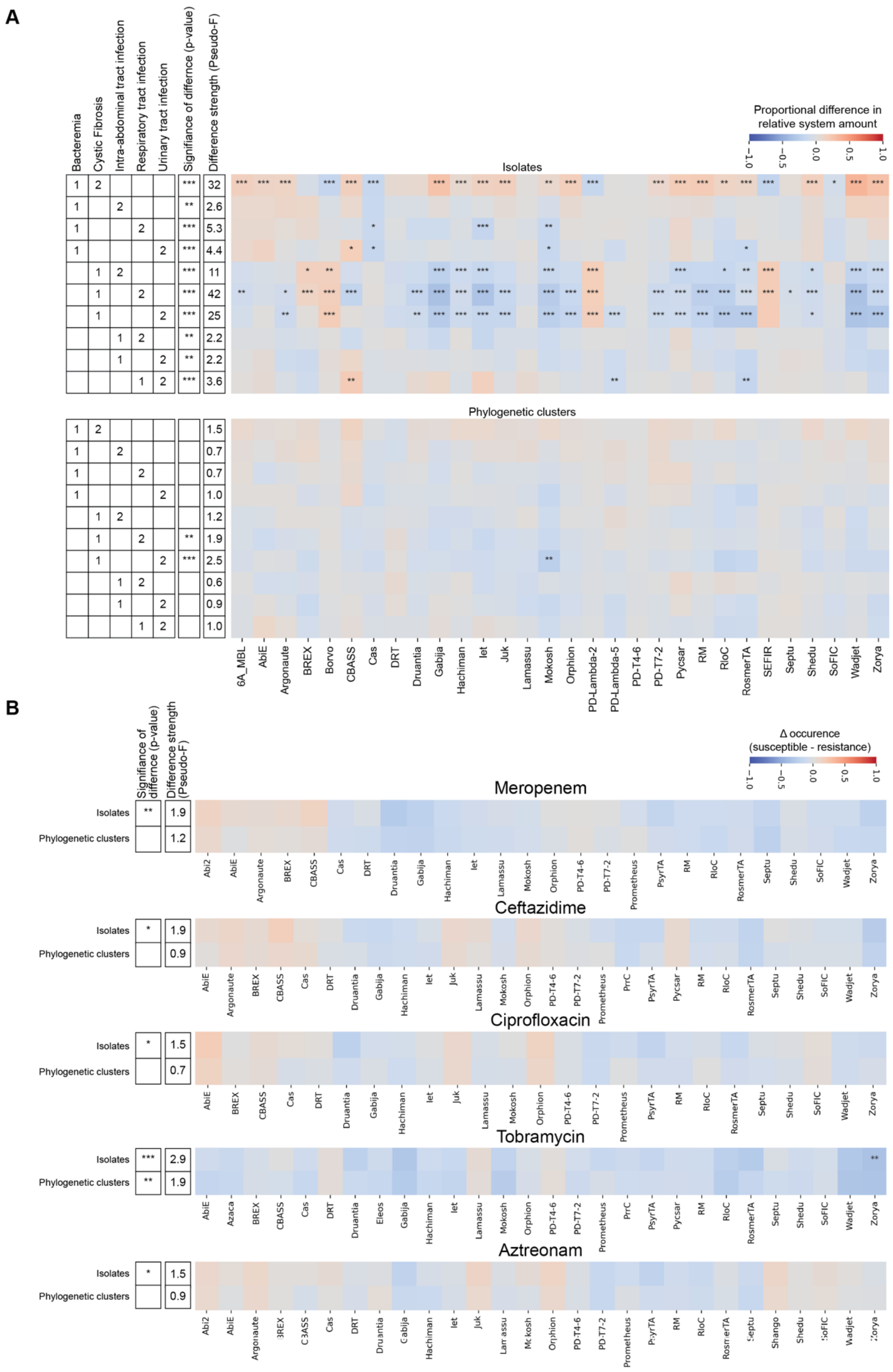


**S5**: Comparisons of the defence system composition for different patient disease indication **(A)** and antibiotic resistances **(B)**. Differences in the total composition were determined using a PERMANOVA test. Heatmap shows the comparisons of individual defence systems between for the different patient disease indication, comparing the proportional difference of how frequently a defence system is found and the statistical significance, based on the p-value of a $\chi^2$-test of independence. Only systems composing a minimum average of 15% of the total defensome in one of the datasets were considered. *** denotes a p-value < 0.0005, ** a p-value < 0.005, * a p-value < 0.05, and n.s. denotes not significant.

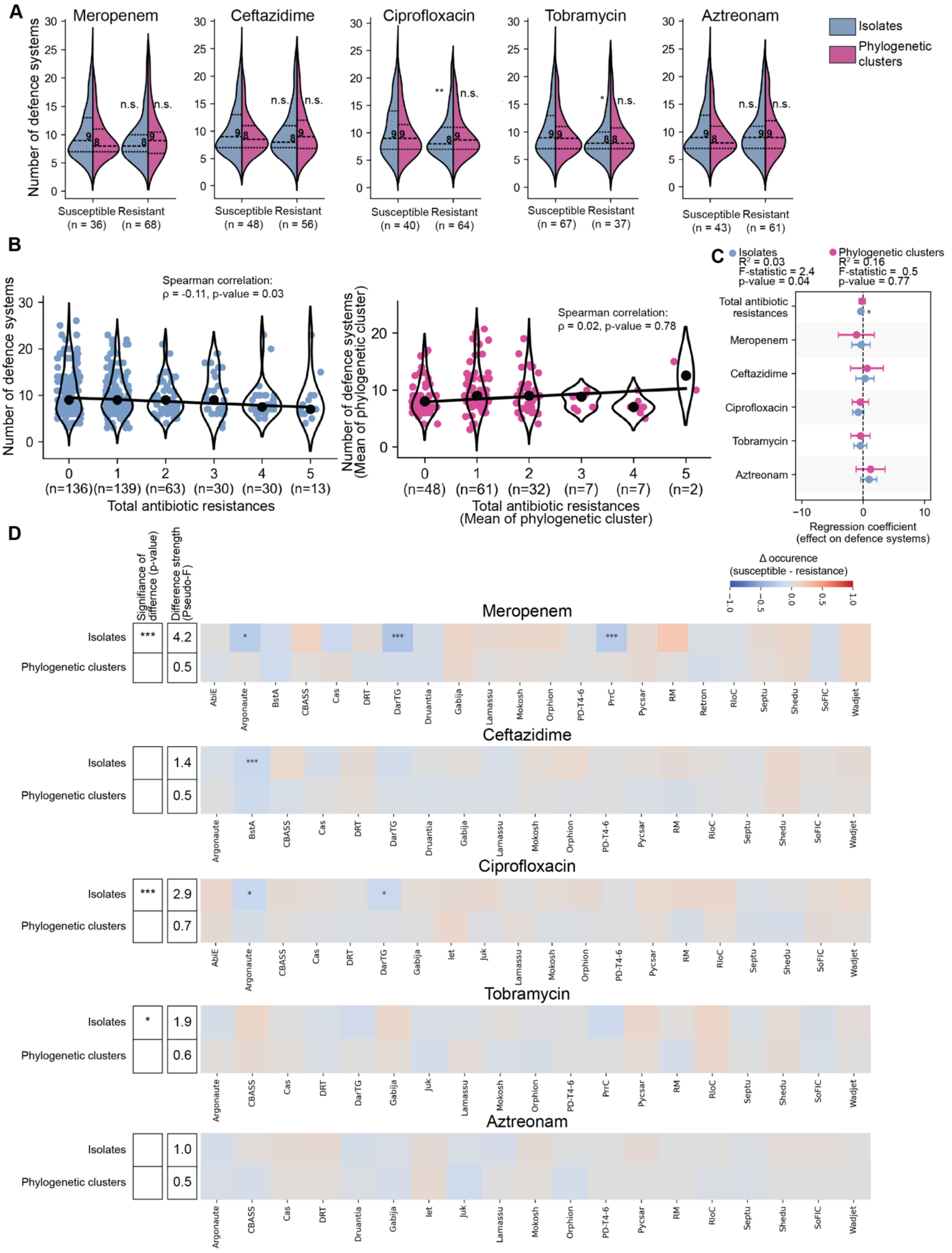


**S6**: Evaluation of the defence systems in *P. aeruginosa* in the Chronic lung database with different antibiotic resistances. **A)** Evaluation of the defence systems in *P. aeruginosa* for antibiotic resistant or susceptible bacteria across all isolates or by averaging each phylogenetic cluster. Five antibiotics from different antibiotic classes are considered. Medians are indicated. Statistical significance of the differences between the databases were calculated using a Welch's T-test for both all isolates (blue) and the phylogenetic clusters (magenta). **B)** Relationship between the number of antibiotics isolates (left) or phylogenetic clusters (right; mean antibiotic resistances used) are resistant to and the average number of defence systems that isolate or phylogenetic cluster (mean number of defence systems used). Violin plots show the distribution of the different average number of defence systems per accumulated antibiotic resistances, individual datapoints are represented in blue (isolates) or magenta (phylogenetic clusters) and the median of each bin is indicated with a black marker. The linear trendline is fitted through the medians of these bins. The Spearman correlation coefficient (ρ) and p-values are reported to indicate the strength and significance of the association, respectively. **C)** Multivariable regression analysis quantifying the association between antibiotic resistance and defence system abundance. Forest plot shows regression coefficients with 95% confidence intervals for multidrug resistance overall and for individual antibiotics. Positive coefficients indicate increased defence system abundance associated with resistance. Models were fitted at both the isolate level (blue) and phylogenetic cluster level (magenta). **D)** Differences in the total composition were determined using a PERMANOVA test. Heatmap shows the comparisons of individual defence systems between for the different antibiotic susceptible and resistant populations, comparing the proportional difference of how frequently a defence system is found and the statistical significance, based on the p-value of a $\chi^2$-test of independence. Only systems composing a minimum average of 15% of the total defensome in one of the datasets were considered. *** denotes a p-value < 0.0005, ** a p-value < 0.005, * a p-value < 0.05, and n.s. denotes not significant.

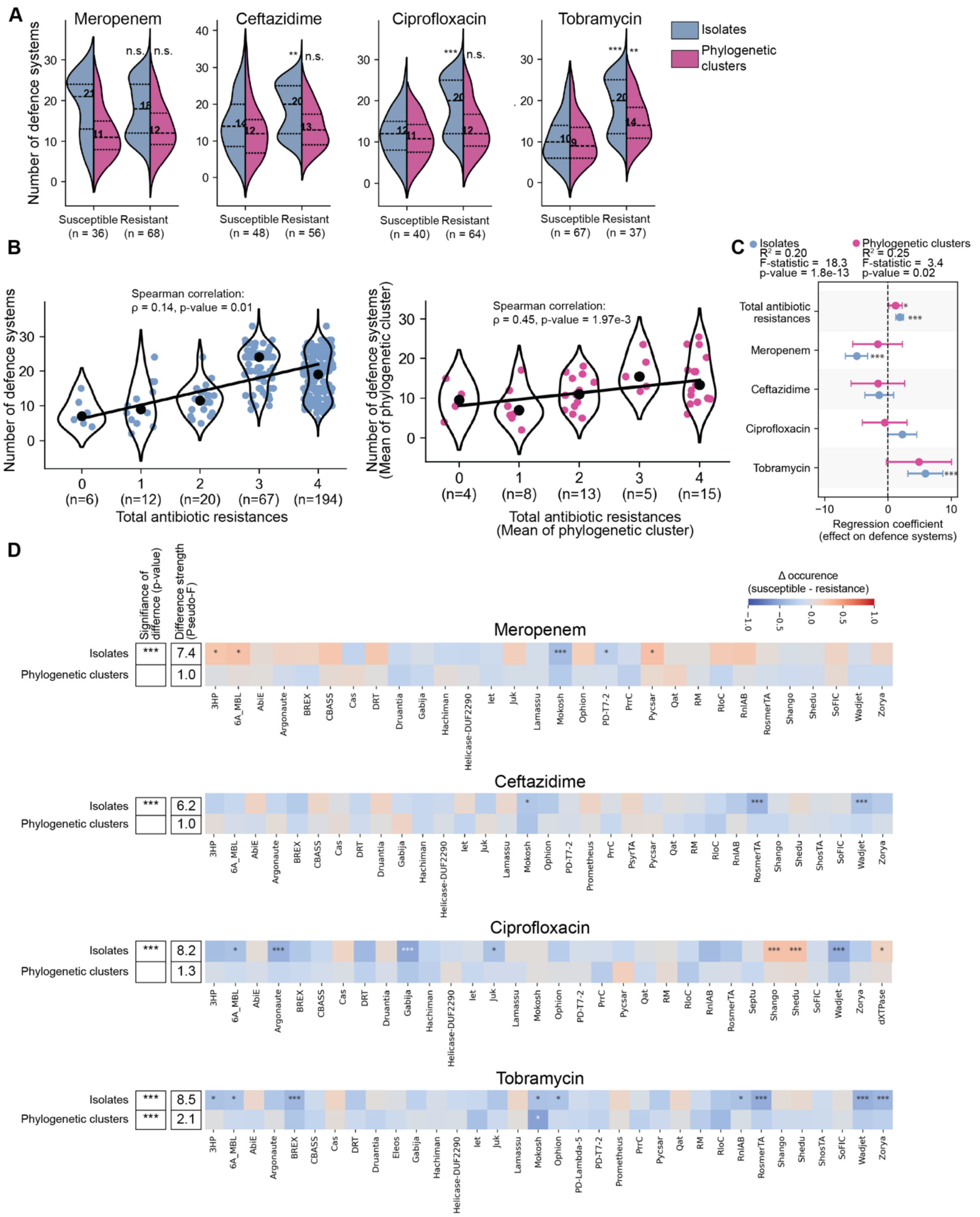

**S7**: Evaluation of the defence systems in *P. aeruginosa* in the Chronic lung database with different antibiotic resistances. **A)** Evaluation of the defence systems in *P. aeruginosa* for antibiotic resistant or susceptible bacteria across all isolates or by averaging each phylogenetic cluster. Five antibiotics from different antibiotic classes are considered. Medians are indicated. Statistical significance of the differences between the databases were calculated using a Welch's T-test for both all isolates (blue) and the phylogenetic clusters (magenta). **B)** Relationship between the number of antibiotics isolates (left) or phylogenetic clusters (right; mean antibiotic resistances used) are resistant to and the average number of defence systems that isolate or phylogenetic cluster (mean number of defence systems used). Violin plots show the distribution of the different average number of defence systems per accumulated antibiotic resistances, individual datapoints are represented in blue (isolates) or magenta (phylogenetic clusters) and the median of each bin is indicated with a black marker. The linear trendline is fitted through the medians of these bins. The Spearman correlation coefficient (ρ) and p-values are reported to indicate the strength and significance of the association, respectively. **C)** Multivariable regression analysis quantifying the association between antibiotic resistance and defence system abundance. Forest plot shows regression coefficients with 95% confidence intervals for multidrug resistance overall and for individual antibiotics. Positive coefficients indicate increased defence system abundance associated with resistance. Models were fitted at both the isolate level (blue) and phylogenetic cluster level (magenta). **D)** Differences in the total composition were determined using a PERMANOVA test. Heatmap shows the comparisons of individual defence systems between for the different antibiotic susceptible and resistant populations, comparing the proportional difference of how frequently a defence system is found and the statistical significance, based on the p-value of a $\chi^2$-test of independence. Only systems composing a minimum average of 15% of the total defensome in one of the datasets were considered. *** denotes a p-value < 0.0005, ** a p-value < 0.005, * a p-value < 0.05, and n.s. denotes not significant.

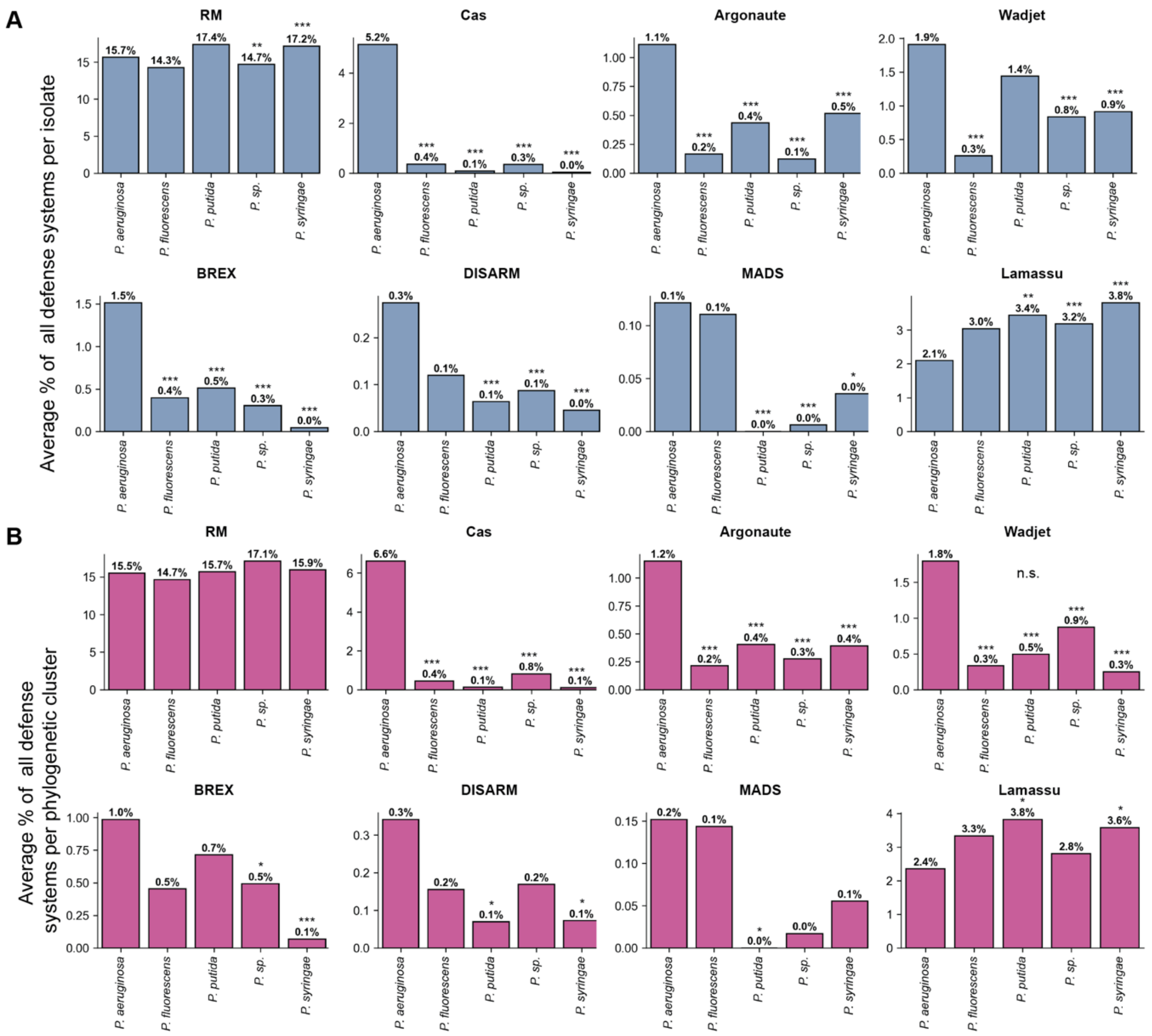


**S8**: Bar graph showing the average contribution of individual anti-plasmid systems to the whole defensome for different *Pseudomonas* species in bacterial isolates **(A)** and phylogenetic clusters **(B)**. Statistical significance of difference compared to *P. aeruginosa* was calculated using a Welch's T-test. *** denotes a p-value < 0.0005, ** a p-value < 0.005, * a p-value < 0.05, and n.s. denotes not significant.

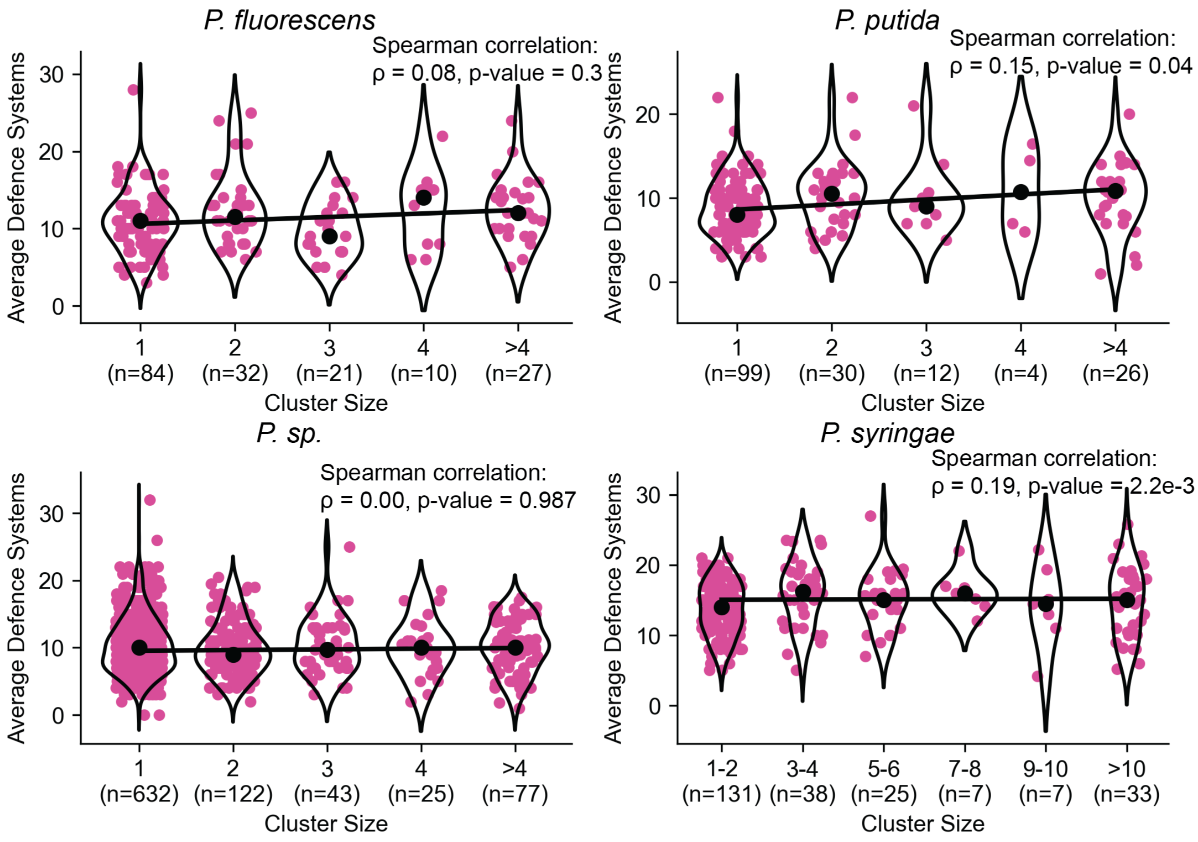


**S9:** Relationship between the number of isolates of phylogenetic clusters and the average number of defence systems per cluster in the Pseudomonas DB for different *Pseudomonas* species. Clusters were grouped in bins based on the number of isolates of that cluster. Violin plots show the distribution of the different average defence systems of a cluster per bin, individual datapoints are represented in magenta and the median of each bin is indicated with a black marker. The linear trendline is fitted through the medians of these bins. The Spearman correlation coefficient (ρ) and p-values are reported to indicate the strength and significance of the association, respectively.
